# Maternal experience-dependent cortical plasticity in mice is circuit- and stimulus-specific and requires MECP2

**DOI:** 10.1101/655035

**Authors:** Billy Y. B. Lau, Keerthi Krishnan, Z. Josh Huang, Stephen D. Shea

## Abstract

The neurodevelopmental disorder Rett syndrome is caused by mutations in the gene *Mecp2*. Misexpression of the protein MECP2 is thought to contribute to neuropathology by causing dysregulation of plasticity. Female heterozygous *Mecp2* mutants (*Mecp2*^*het*^) failed to acquire a learned maternal retrieval behavior when exposed to pups, an effect linked to disruption of parvalbumin-expressing inhibitory interneurons (PV+) in the auditory cortex. However, the consequences of dysregulated PV+ networks during early maternal experience for auditory cortical sensory activity are unknown. Here we show that maternal experience in wild-type adult female mice (*Mecp2*^*wt*^) triggers suppression of PV+ auditory responses. We also observe concomitant disinhibition of auditory responses in deep-layer pyramidal neurons that is selective for behaviorally-relevant pup vocalizations. These neurons also exhibit sharpened tuning for pup vocalizations following maternal experience. All of these neuronal changes are abolished in *Mecp2*^*het*^, yet a genetic manipulation of GABAergic networks that restores accurate retrieval behavior in *Mecp2*^*het*^ also restores maternal experience-dependent plasticity of PV+. Our data are consistent with a growing body of evidence that cortical networks are particularly vulnerable to mutations of *Mecp2* in PV+ neurons.

## INTRODUCTION

In humans, loss of function mutations in the X chromosome-linked gene *MECP2* cause the neurodevelopmental disorder Rett Syndrome (RTT) (Amir et al., 1999). MECP2, the protein product of *MECP2*, binds to methylated DNA (Lewis et al., 1992) and controls transcriptional programs by modifying chromatin structure (Skene et al., 2010; Agarwal et al., 2011; Brink et al., 2013; Maxwell et al., 2013). The vast majority of humans with RTT are females with a heterozygous loss of function mutation who are therefore mosaic for the gene (Van den Veyver and Zoghbi, 2000). Mounting evidence from mice that carry mutations abolishing expression of MECP2 strongly suggests that it is an important regulator of plasticity in development and adulthood (Deng et al., 2010; McGraw et al., 2011; Noutel et al., 2011; Na et al., 2013; Deng et al., 2014; He et al., 2014; Krishnan et al., 2015; Tai et al., 2016; Krishnan et al., 2017; Gulmez Karaca et al., 2018). Nevertheless, little is known about how these mutations affect *in vivo* sensory activity (Durand et al., 2012; Banerjee et al., 2016) and whether this has direct consequences for learning and behavior.

MECP2 is expressed in all cells in the brain, yet parvalbumin-expressing cortical inhibitory interneurons (PV+) appear to be especially vulnerable to loss of MECP2 (Ito-Ishida et al., 2015; He et al., 2017; Morello et al., 2018). Selective knockout of *Mecp2* in either PV+ or forebrain inhibitory neurons broadly recapitulates major aspects of the phenotype seen in whole mouse knockouts (Chao et al., 2010; Ito-Ishida et al., 2015). Parvalbumin is a calcium binding protein that marks a subtype of cortical interneurons that share functional properties. Among other functions (Cardin et al., 2009; Sohal et al., 2009; Atallah et al., 2012; Lee et al., 2012; Wilson et al., 2012; Kim et al., 2016), PV+ play a crucial role in regulating learning and plasticity during development and adulthood (Donato et al., 2013). Specifically, modulation of PV+ networks controls the opening and closing of windows of heightened plasticity, such as during developmental critical periods (Levelt and Hubener, 2012; Takesian and Hensch, 2013). Therefore, dysregulation of PV+ in *Mecp2* mutant mice disrupts developmental critical periods and adult learning (Picard and Fagiolini, 2019).

For example, we recently showed that *Mecp2* mutations impair auditory cortical plasticity and maternal vocal perception in adult female mice through circuitry changes that involve PV+ (Krishnan et al., 2017). Primiparous female mice, or virgin females exposed to pups (surrogates), both exhibit experience-dependent improvement in retrieving isolated pups in response to their ultrasonic distress cries (Sewell, 1970; Ehret et al., 1987). Moreover, exposure of naïve virgin wild-type females to pups also triggers changes to inhibitory networks in the auditory cortex that affect neural coding of pup vocalizations (Liu and Schreiner, 2007; Galindo-Leon et al., 2009; Cohen et al., 2011; Lin et al., 2013; Cohen and Mizrahi, 2015; Marlin et al., 2015). Interestingly, we found that additional changes occur specifically to PV+ in the auditory cortex of heterozygous *Mecp2* mutants (*Mecp2*^*het*^), namely increased expression of parvalbumin and intensification of perineuronal nets (PNNs). PNNs are glycoprotein structures in the extracellular matrix surrounding PV+ that act as physical barriers to or modulators of synaptic modifications, and actively control maturation of PV+ (Krishnan et al., 2015; Sorg et al., 2016; Miyata and Kitagawa, 2017; Sigal et al., 2019). Thus, PNNs are proposed to work in concert with elevated parvalbumin to act as ‘brakes’ on plasticity e.g. at the termination of developmental critical periods (Takesian and Hensch, 2013). In support of this conclusion, genetic reduction of levels of the GABA-synthesizing enzyme GAD67 in the *Mecp2*^*het*^ background normalized PV and PNN expression and restored maternal learning (Krishnan et al., 2017). Taken together, these findings predict that changes in sensory responses to pup vocal signals in wild-type females are reshaped by PV+ following maternal experience, and that this sensory plasticity is disrupted in *Mecp2*^*het*^.

Here we use single-neuron electrophysiological recordings from awake, head-fixed mice combined with spike waveform analysis to show that maternal experience in wild-type females (*Mecp2*^*wt*^) triggers reduced inhibition and increased excitation in the auditory responses of presumptive deep-layer pyramidal neurons. Direct recordings from putative PV+ neurons show decreased stimulus-evoked spiking following maternal experience, suggesting that the changes in the pyramidal cells may reflect disinhibition by the PV+ neurons. Changes in the auditory responses of deep-layer pyramidal neurons are selective for behaviorally-relevant pup vocalizations, as opposed to synthetic stimuli such as pure tones. The deep-layer pyramidal neurons in *Mecp2*^*wt*^ also exhibit sharpened tuning for pup vocalizations following maternal experience. None of these neuronal changes were observed in *Mecp2*^*het*^. In fact, in these mice we found that PV neuron output was actually increased in surrogate. Notably, the same GAD67 perturbation that restored retrieval behavior also reinstated PV neuron plasticity in response to maternal experience.

Considering these findings together with our previous results, we propose that MECP2 acts in cortical PV+ interneurons to coordinate stimulus-specific, experience-dependent cortical plasticity, thereby reshaping the output of deep layer pyramidal neurons. We propose that mutations in *Mecp2* disrupt adult plasticity of cortical representations by arresting modulation of cortical PV+ networks, and that this has negative behavioral consequences.

## RESULTS

Our previous work revealed molecular expression changes to inhibitory networks in the auditory cortex of surrogate female mice (Sur) that were co-housed with a mother and her pups beginning at their birth, as compared to maternally-inexperienced virgin females (Naive). We further showed that the changes to inhibitory circuitry were significantly altered in *Mecp2*^*het*^, and that independent genetic and pharmacological interventions that partially restored wild type marker expression significantly improved behavioral performance (Krishnan et al., 2017). We speculated that the molecular expression differences between surrogate *Mecp2*^*wt*^ (SurWT) and *Mecp2*^*het*^ (SurHET) might lead to or accompany differences in sensory activity in the auditory cortex. Therefore, we performed loose patch extracellular recordings from individual neurons in the auditory cortex of awake, head-fixed female mice of both genotypes, before and after maternal experience.

### Photoidentification of PV+ neurons

Previous studies have established that PV+ cortical neurons can be identified in electrophysiological recordings by the characteristics of their extracellular waveform (Wu et al., 2008; Oswald and Reyes, 2011; Cohen and Mizrahi, 2015). In one particularly relevant example, Cohen and Mizrahi (2015) made extracellular recordings from auditory cortical neurons under guidance by two photon laser scanning microscopy. They reported that genetically labeled PV+ formed a cluster distinct from their PV-pyramidal cell neighbors on a scatterplot of 1) the time between the positive peak of the spike waveform and the negative trough, and 2) the ratio of their amplitude.

We confirmed this finding in our recordings, using photoidentification (Lima et al., 2009) of PV+ neurons in a mouse line expressing Channelrhodopsin2 only in PV+ neurons (Figure 1A). PV+ neurons were identified in recordings by their spiking response to blue light (473 nm), delivered through a light fiber placed next to the recording pipette. ChR2-expressing PV+ (n = 9) were unambiguously identified by their highly reliable (> 80% of trials) and short median latency (< 5 ms) response to light (Figure 1B, C). Other cells recorded during the same experiments (n = 17) fired rarely to light (< 25% of trials) and with very long median latencies (> 50 ms), and they were deemed non-light responsive, and therefore likely PV-(Figure 1B, C).

**Figure 1:**
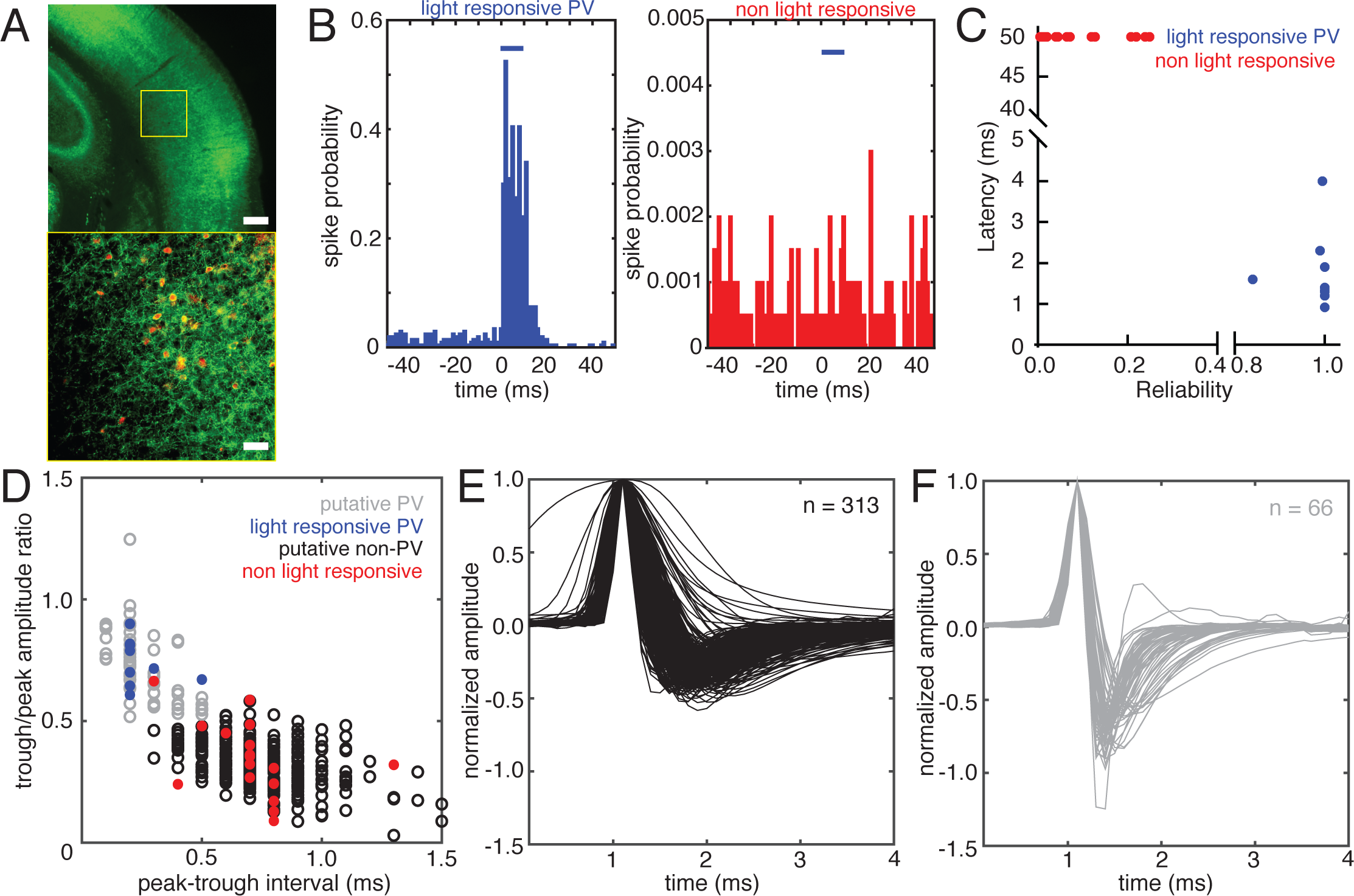
Optogenetic identification of PV neurons during *in vivo* neuronal recordings from auditory cortex in awake, head-fixed mice. (A) Photomicrographs taken at 4x (*upper panel*) and 20x (*lower panel*) of a section of the auditory cortex in an *Ai32;PV-ires-Cre* mouse expressing ChR2 in PV neurons. ChR2 was fused to GFP and is visualized in green. The yellow box in the upper panel denotes the location of the image in the lower panel. PV is visualized in red. Scale bars are 200 μm (upper) and 50 μm (lower). (B) Representative example histograms of responses in a positively identified PV neuron *(left)* and a presumed non-PV neuron *(right)* to 10 ms light pulses (blue bars, 473 nm) Spike probability is computed as the number of spikes in each bin divided by the number of light trials. (C) Scatterplot of the reliability of light responses (fraction of trials exhibiting spikes in response to light within the first 50 ms after light onset) versus the latency of spikes in response to light (n = 26). ChR2-expressing PV neurons (n = 9) are distinguished from non-ChR2-expressing presumed non-PV neurons (n = 17) by high reliability and low latency responses. (D) Scatterplot of spike waveform features for all neurons reported in this study (n = 379). The quantities plotted are the time in ms between the peak and the trough of the averaged waveform (see Methods) versus the ratio of the amplitudes of the trough and the peak of the averaged waveform. Blue and red points denote the light-responsive and light non-responsive neurons shown in (C), respectively. Based on their locations in the scatter plot relative to the larger waveform dataset, the remaining neuronal waveforms were classified as putative PV neurons (gray, n = 66) and putative non-PV neurons (black, n = 313). (E, F) Plots of the averaged spike waveform from all putative non-PV (E) and putative PV (F) neurons recorded for this study. Each waveform is aligned and normalized to its peak amplitude.

Figure 1D depicts a scatterplot of the peak-trough interval in milliseconds versus the trough/peak amplitude ratio for all *Mecp2*^*wt*^ and *Mecp2*^*het*^ cells reported in this study (n = 379). The blue dots denote data points for photo-identified PV+, and the red dots denote data points for cells that did not respond to light. Based on the segregation of optically-identified PV+ to the quadrant of the plot bounded by peak-trough intervals < 0.6 ms and trough/peak amplitude ratios > 0.5, we designated all cells in that quadrant as putative PV+ (n = 66) (Figure 1F). All other cells were designated as putative non-PV (n = 313) (Figure 1E), and likely are primarily pyramidal neurons (Cohen and Mizrahi, 2015). For simplicity and clarity, we henceforth refer to these putatively-identified populations as ‘PV’ and ‘non-PV.’

As is typically observed, we found that PV neurons had higher spontaneous firing rates than non-PV cells across both genotypes and behavioral conditions (Figure 2A; non-PV: 3.35 [3.0 3.8] spikes/s; PV: 7.50 [6.2 8.7] spikes/s, Mann-Whitney U test; *p* < 0.001) (all values reported as median and [95% confidence interval] throughout). Across behavioral conditions, non-PV neurons showed significantly lower spontaneous firing rates in *Mecp2*^*het*^ as compared to *Mecp2*^*wt*^ (Figure 2B; WT: 3.96 [3.4 4.9] spikes/s; HET: 2.60 [2.0 3.3] spikes/s; Mann-Whitney U test, *p* < 0.001). Spontaneous firing rates did not differ between PV+ neurons in *Mecp2*^*wt*^ and *Mecp2*^*het*^. (Figure 2C; WT: 7.53 [5.2 11] spikes/s; HET: 7.48 [5.8 8.6] spikes/s; Mann-Whitney U test, *p* = 0.54). Spontaneous activity in neither cell type and neither genotype was affected by maternal experience (p > 0.05).

**Figure 2:**
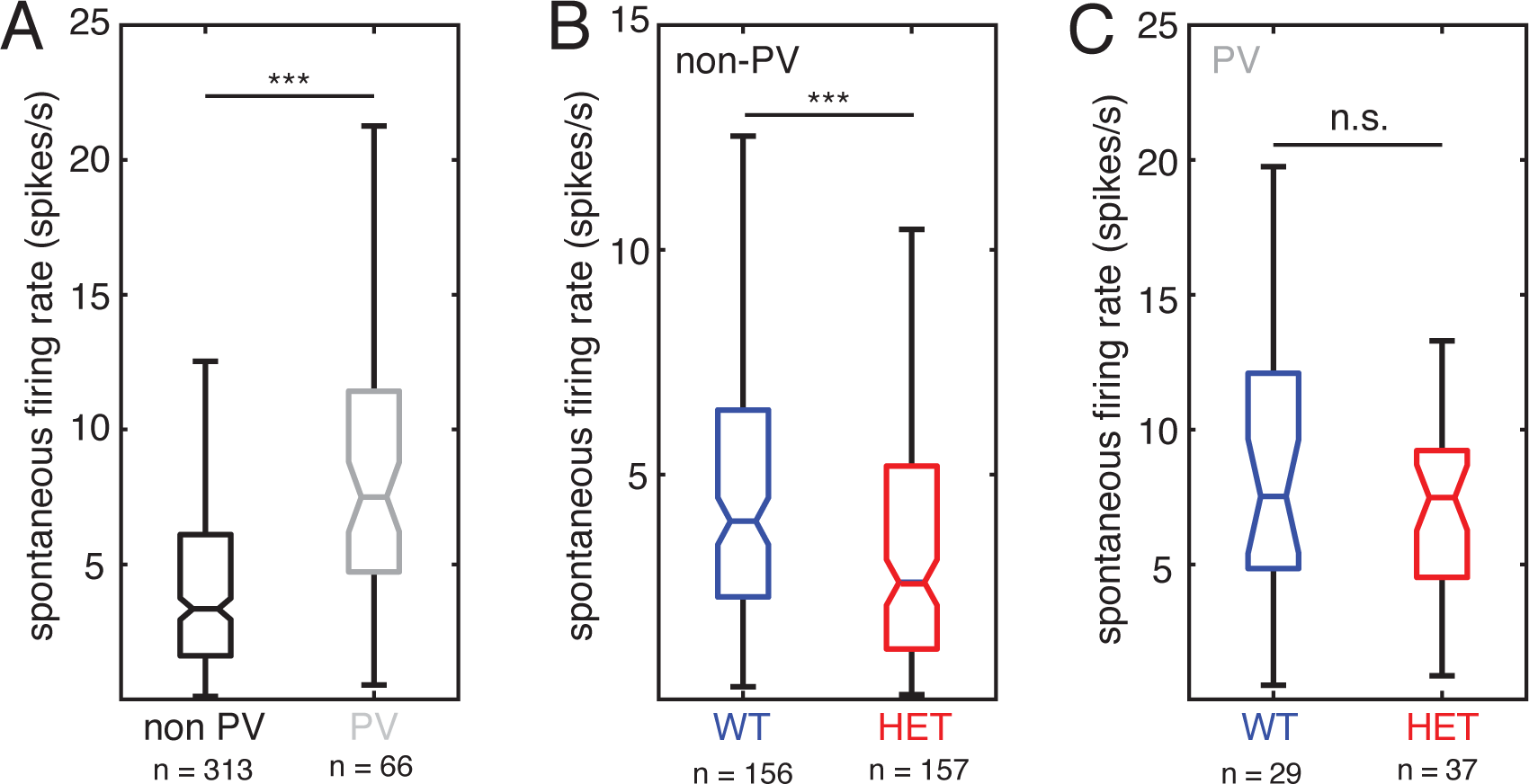
Spontaneous firing rates differ between cell-types and genotypes. (A) Boxplot of spontaneous firing rates comparing all non-PV neurons (black) and all PV neurons (gray) (Mann-Whitney U test; *\*\*p* < 0.001). (B) Boxplot of spontaneous firing rates comparing all non-PV neurons from WT females (blue) and all non-PV neurons from HET females (red) (Mann-Whitney U test; *\*\*p* < 0.001). (C) Boxplot of spontaneous firing rates comparing all PV neurons from WT females (blue) and all PV neurons from HET females (red) (Mann-Whitney U test; n.s. *p* = 0.54).

### Responses to pup calls are strengthened in SurWT, but not SurHET

We recorded auditory responses in the left ‘core’ (thalamorecipient) auditory cortex (Shepard et al., 2016) evoked by a library of eight different pup call exemplars recorded from a range of ages in our laboratory (Figure 3A). We recorded from 18 *Mecp2*^*wt*^ (14 naïve and 4 surrogates) and 22 *Mecp2*^*het*^ (13 naïve and 9 surrogates) for a total of 313 putative non-PV neurons and 66 putative PV neurons. Individual neurons exhibited distinct responses to each exemplar. As a typical example, Figure 3B shows raster plots and peristimulus time histograms (PSTHs) for each call compiled from a recording of a single non-PV neuron. Some calls evoked firing rate suppression (‘inhibition’), while others evoked firing rate increases (‘excitation’). PV neurons also typically exhibited distinct changes in firing rate to each call (Figure 3C).

**Figure 3:**
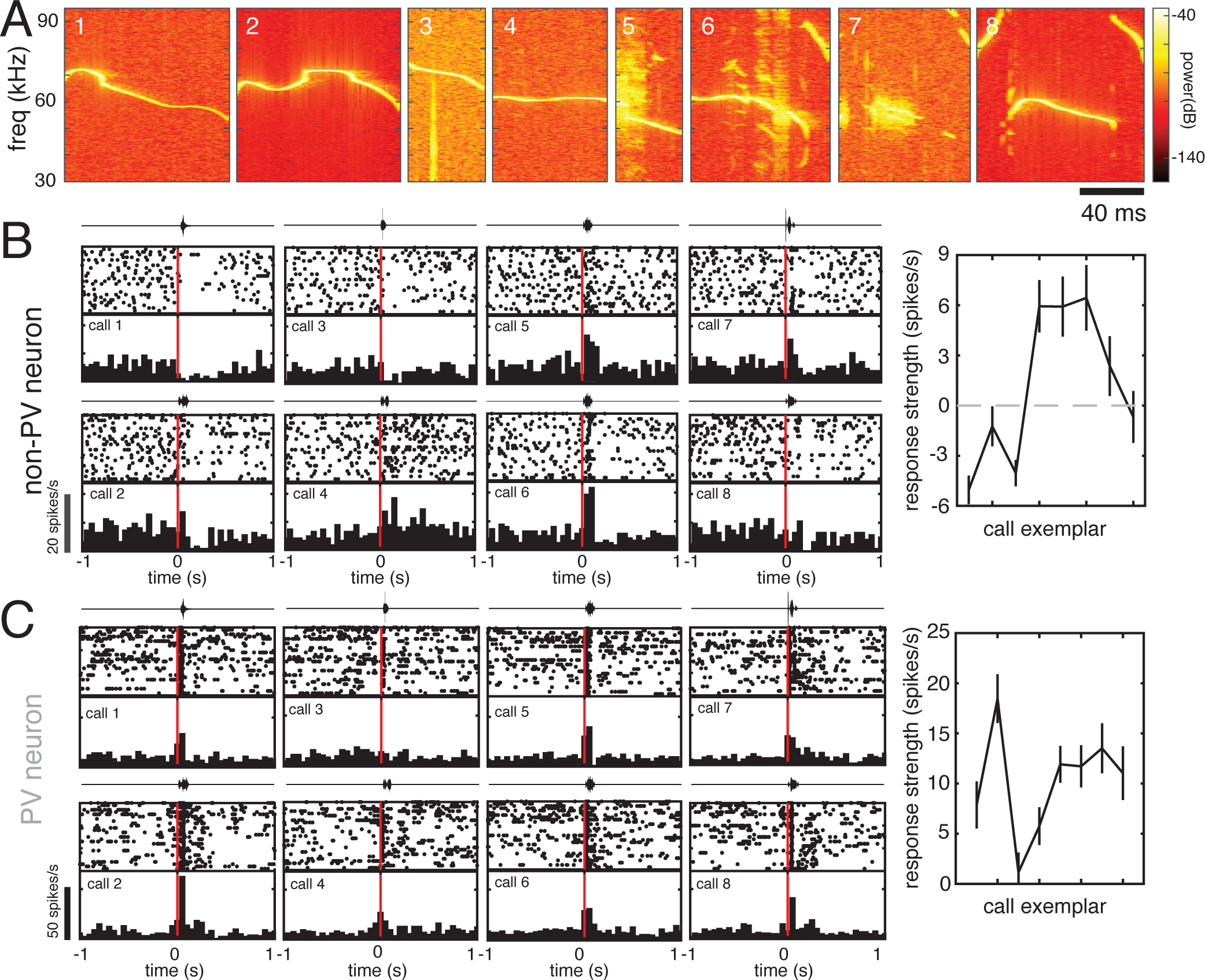
Individual auditory cortical neurons exhibit diverse responses to different pup call exemplars. (A) Spectrograms of all call stimuli used in this study. (B, C) Auditory responses of example non-PV (B) and PV (C) neurons from naive WT females to a set of eight different pup call exemplars. Diverse responses to each of eight calls are depicted by a raster plot and peristimulus time histogram (PSTH) (20 trials/call; bin size = 50 ms). Above each raster and PSTH is the oscillograph of the call played during those trials. On the right is a tuning curve plotting the mean change in firing rate during the first 200 ms after call onset (response strength). The use of the term “tuning curve” is in this case not intended to imply systematic neighbor relationships between stimuli on the x-axis.

As an initial assessment of maternal experience-dependent changes in neural encoding of pup calls, we examined the changes in median stimulus-evoked firing rates with maternal experience in each genotype. In *Mecp2*^*wt*^, the median response strength (baseline-subtracted firing rate; see Methods) of non-PV cells to playback of pup calls was higher in SurWT than in NaiveWT (Figure 4A; NaiveWT: −0.28 [-0.5 0.05] spikes/s; SurWT: 0.73 [0.1 1.0] spikes/s; Mann-Whitney U test, *p* < 0.001). In contrast, SurHET showed a significant decrease in median pup call response strength as compared to NaiveHET (Figure 4A; NaiveHET: 0.37 [0.1 0.7] spikes/s; SurHET: −0.018 [-0.2 0.1] spikes/s, Mann-Whitney U test; *p* < 0.01).

**Figure 4:**
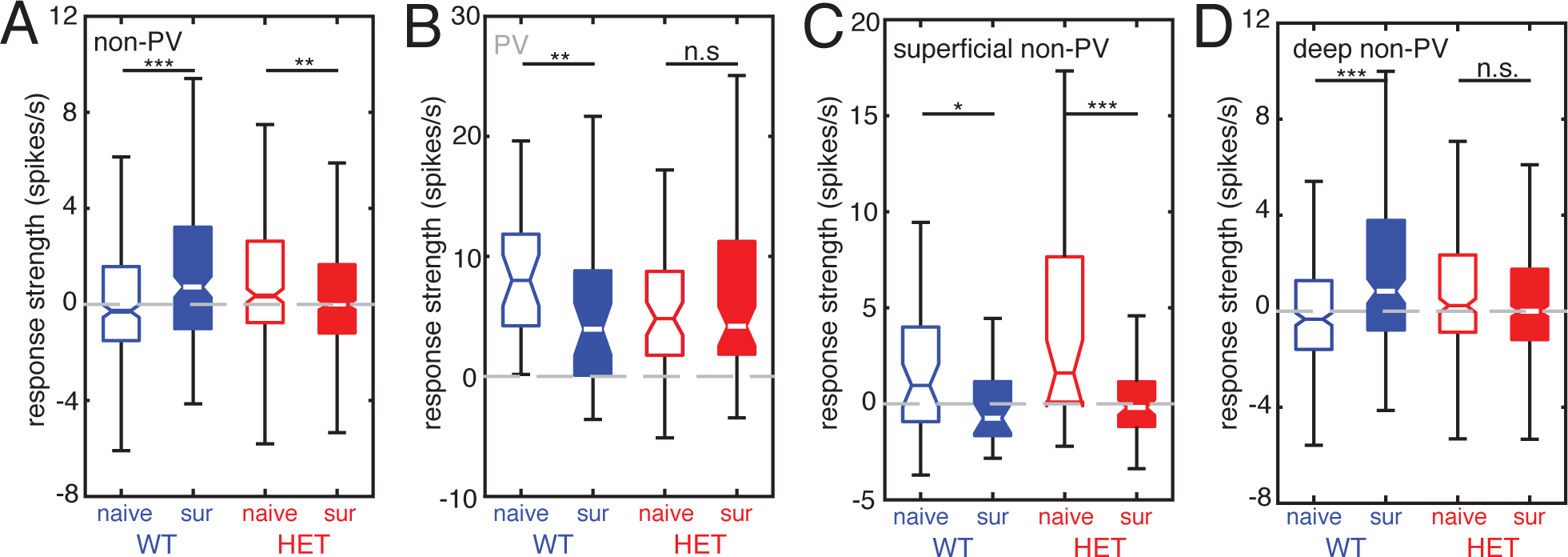
Maternal experience triggers distinct cell type- and layer-specific changes to pup call responses in WT and HET females. (A) Boxplot of mean response strength to all calls for all non-PV cells comparing naive females to surrogate females for WT (blue; naive n = 344 cell-call pairs and surrogate n = 224 cell-call pairs) and HET (red; naive n = 344 cell-call pairs and surrogate n = 456 cell-call pairs) genotypes (Mann-Whitney U test with Holm-Bonferroni correction; *\*\*\*p* < 0.001; *\*\*p* < 0.01). (B) Boxplot of mean response strength to all calls for all PV cells comparing naive females to surrogate females for WT (blue; naive n = 32 cell-call pairs and surrogate n = 40 cell-call pairs) and HET (red; naive n = 64 cell-call pairs and surrogate n = 80 cell-call pairs) genotypes (Mann-Whitney U test with Holm-Bonferroni correction; \**\*p* < 0.01; n.s. = not significant, *p* = 0.87). (C) Boxplot of mean response strength to all calls for all superficial (depth < 500 μm) non-PV cells comparing naive females to surrogate females for WT (blue; naive n = 48 cell-call pairs and surrogate n = 32 cell-call pairs) and HET (red; naive n = 48 cell-call pairs and surrogate n = 80 cell-call pairs) genotypes (Mann-Whitney U test with Holm-Bonferroni correction; *\*p* < 0.05; *\*\*\*p* < 0.001). (D) Boxplot of mean response strength to all calls for all deep (depth < 500 μm) non-PV cells comparing naïve females to surrogate females for WT (blue; naive n = 296 cell-call pairs and surrogate n = 192 cell-call pairs) and HET (red; naive n = 296 cell-call pairs and surrogate n = 376 cell-call pairs) genotypes (Mann-Whitney U test with Holm-Bonferroni correction; *\*\*\*p* < 0.001; n.s. *p* = 0.07).

Our previous work (Krishnan et al., 2017) indicated that maternal experience triggers changes to inhibitory networks. Nevertheless, these changes in firing rate in response to call playback could, in principle, be due to either changes in excitatory or inhibitory input. Recordings from inhibitory PV neurons suggest that the increase in auditory responses of non-PV neurons in SurWT may be due to reduced inhibitory input from PV neurons. Specifically, in *Mecp2*^*wt*^, increased call responses in non-PV cells of SurWT over NaiveWT was mirrored by significantly reduced call responses in PV neurons (Figure 4B; NaiveWT: 8.00 [4.5 12] spikes/s; SurWT: 3.94 [1.2 7.3] spikes/s; Mann-Whitney U test; *p* < 0.01). On the other hand, no change in response strength was seen between NaiveHET and SurHET PV neurons (Figure 4B; NaiveHET: 4.82 [3.3 6.0] spikes/s; SurHET: 4.19 [3.6 7.0] spikes/s; Mann-Whitney U test, *p* = 0.87).

We next assessed whether the increase in pup call responses after maternal experience was layer-specific. We therefore sorted our recordings from non-PV cells according to depth, designating cells that were recorded <500 μm from the surface as superficial non-PV neurons and cells that were recorded at 500 – 1100 μm from the surface as deep non-PV neurons. Both *Mecp2*^*wt*^ and *Mecp2*^*het*^ showed significant decreases in the median response strength of superficial non-PV neurons to pup calls following maternal experience (Figure 4C; NaiveWT: 0.96 [-0.7 3] spikes/s; SurWT: −0.75 [-2 1] spikes/s; Mann-Whitney U test, *p* < 0.05; NaiveHET: 1.61 [0.57 3.9] spikes/s; SurHET: −0.19 [-0.8 0.4] spikes/s; Mann-Whitney U test, *p* = 0.001). However, increased pup call responses were observed in deep layer non-PV in *Mecp2*^*wt*^ (Figure 4D; NaiveWT: −0.33 [-0.6 −0.1] spikes/s; SurWT: 0.83 [0.4 1.6] spikes/s; Mann-Whitney U test, *p* < 0.001), but not in *Mecp2*^*het*^ (Figure 4D; NaiveHET: 0.23 [-0.05 0.6] spikes/s; SurHET: 0.00 [-0.2 0.2] spikes/s; Mann-Whitney U test, *p* = 0.07). Taken together, these results suggest that in *Mecp2*^*wt*^, auditory responses to pup calls, specifically in deep layer non-PV neurons increase following maternal experience due to reduced inhibition by PV neurons. It is worth noting that of the 27 call-responsive PV neurons reported here, all but four were located in the deep layers. Considered in light of our previous work functionally implicating PV neurons in vocal perception behavior, these data also raise the possibility that plasticity in deep non-PV cells may be important for maternal behavioral learning, since it is disrupted in *Mecp2*^*het*^.

### Plasticity of auditory responses in non-PV neurons is stimulus-specific

Auditory responses of non-PV neurons to a sequence of synthetic and behaviorally-irrelevant tone and noise stimuli were not affected by maternal experience in *Mecp2*^*wt*^. Median response strength to synthetic stimuli did not differ in these neurons between NaiveWT and SurWT (Figure 5A; NaiveWT: −2.25 [2.6 1.8] spikes/s; SurWT: −2.23 [-2.6 −2.0] spikes/s; Mann-Whitney U test, *p* = 0.84). PV neuron responses to these stimuli were still significantly reduced in surrogates (Figure 5B; NaiveWT: 0.21 [-3.0 2.9] spikes/s; SurWT: −3.7 [-6.3 −2.1] spikes/s; Mann-Whitney U test, *p* < 0.01), but notably, the responses in non-PV were not affected. Potentially, this reflects a greater influence of PV-mediated inhibition on non-PV neuron responses to pup vocalizations as compared to synthetic sounds.

**Figure 5:**
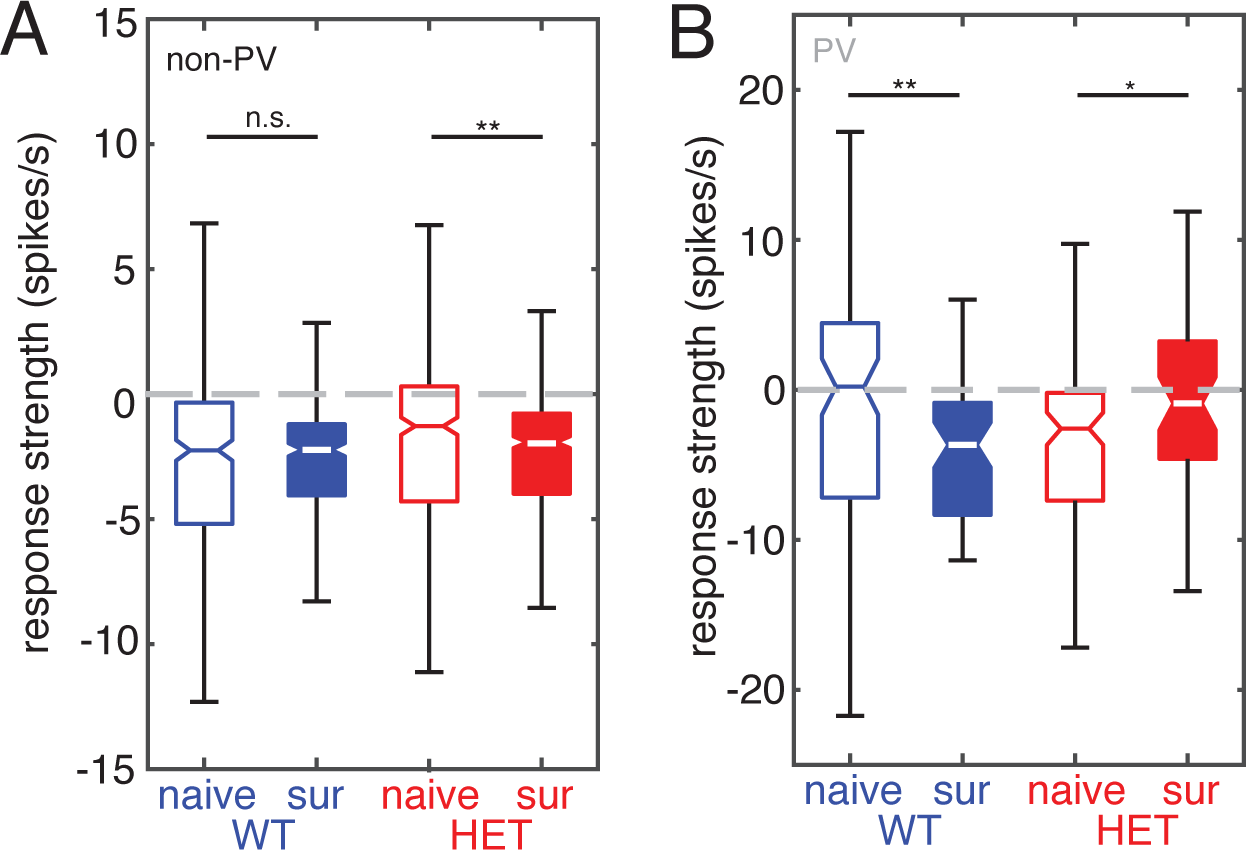
Maternal experience-dependent plasticity in WT females is stimulus-specific. (A) Boxplot of mean response strength to tones and noise stimuli for all non-PV cells comparing naive females to surrogate females for WT (blue; naive n = 342 cell-stimulus pairs and surrogate n = 288 cell-stimulus pairs) and HET (red; naive n = 432 cell-stimulus pairs and surrogate n = 462 cell-stimulus pairs) genotypes (Mann-Whitney U test with Holm-Bonferroni correction; n.s. *p* = 0.84; *\*\*p* < 0.01). (B) Boxplot of mean response strength to tones and noise stimuli for all PV cells comparing naive females to surrogate females for WT (blue; naive n = 96 cell-stimulus pairs and surrogate n = 60 cell-stimulus pairs) and HET (red; naive n = 108 cell-stimulus pairs and surrogate n = 48 cell-stimulus pairs) genotypes (Mann-Whitney U test with Holm-Bonferroni correction; *\*\*p* < 0.01; \**p* < 0.05).

In *Mecp2*^*het*^, responses to the synthetic stimuli were affected by maternal experience in the same manner as responses to pup calls. Specifically, non-PV neurons in SurHET showed slightly, but significantly lower firing rates than NaiveHET (Figure 5A; NaiveHET: −1.28 [-1.8 −1.1] spikes/s; SurHET: −1.92 [-2.2 −1.7] spikes/s; Mann-Whitney U test, *p* < 0.01). In contrast to wild type mice, median response strength of PV neurons to synthetic stimuli was slightly but significantly increased between NaiveHET and SurHET (Figure 5B; NaiveHET: −2.58 [-4.5 −1.3] spikes/s; SurHET: −0.90 [-3.5 1.3] spikes/s; Mann-Whitney U test, *p* < 0.05).

### Plasticity of auditory responses in deep non-PV neurons affects the late response to calls

We next analyzed the temporal structure of inhibitory and excitatory responses separately. In wild type mice, many responses in both PV and non-PV neurons exhibited biphasic structure, with an early component and a late component that appeared to be independently regulated by maternal experience. Stimulus onset and offset responses have been widely observed in auditory cortical neurons and are likely to arise from distinct synaptic inputs (Scholl et al., 2010). We found that in *Mecp2*^*wt*^, in addition to an overall shift of responses in favor of excitation, response plasticity specifically in the later part of the response effectively extended the excitatory responses of deep non-PV neurons.

First, we identified all cell-call pairs (mean response of one cell to one specific call) from deep layer non-PV neurons that resulted in a significant change in firing rate using a bootstrap test (see Methods). Then, the bins of all response PSTHs for each cell were normalized to a Z score. Figure 6A (upper panels) depicts two-dimensional PSTHs of all significant responses for each combination of genotype and maternal experience condition sorted from the most inhibitory responses to the most excitatory responses. The traces below (Figure 6A) represent the mean of all net excitatory and all net inhibitory responses comparing the magnitude and temporal structure between naive mice (black) and surrogate mice (red).

**Figure 6:**
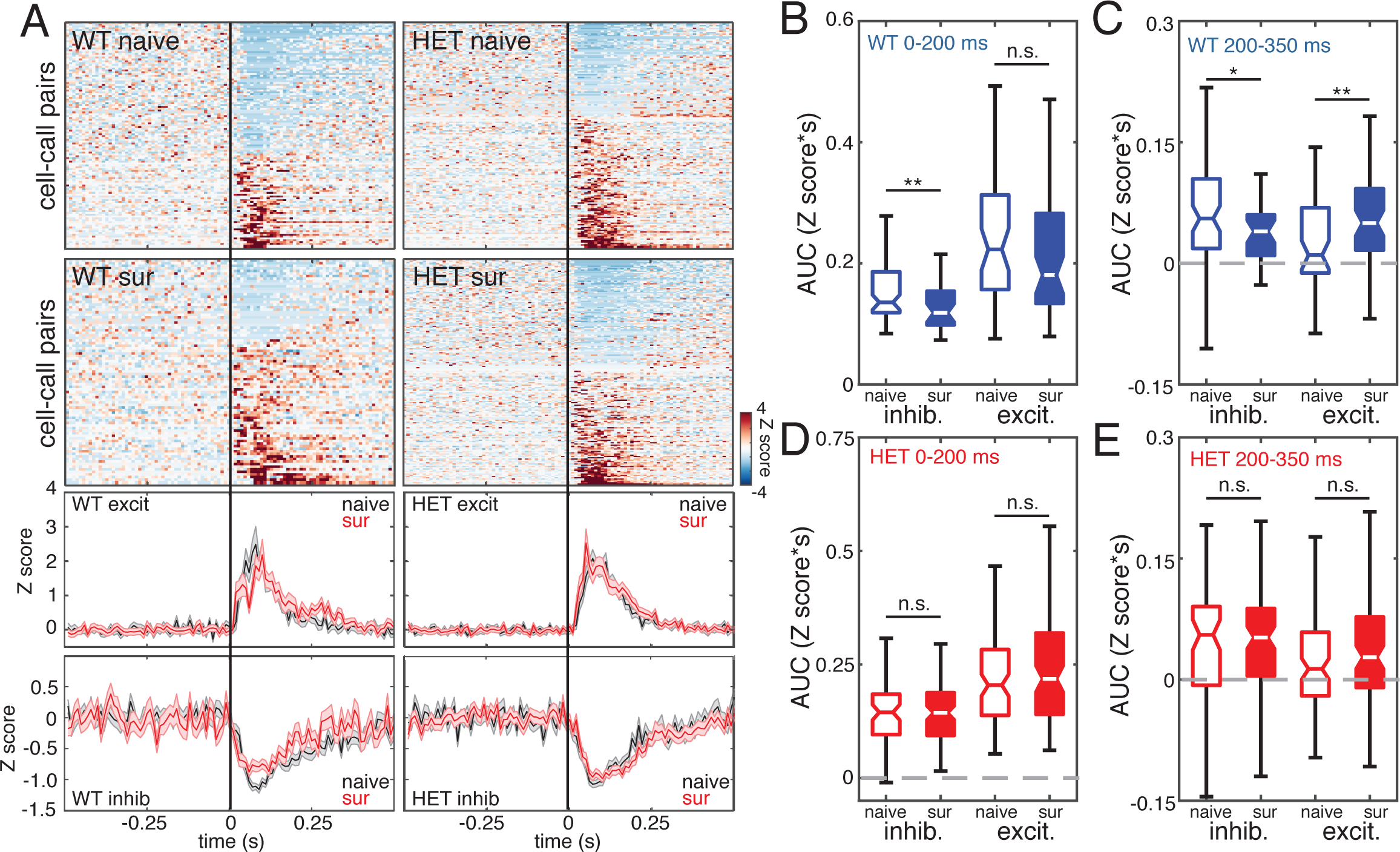
Excitatory responses of deep non-PV neurons to pup calls are disinhibited and extended in duration by maternal experience in WT females. (A) *Top panels:* Heat maps depicting Z score-normalized responses for all deep non-PV cell-call pairs in naive and surrogate WT females and in naive and surrogate HET females. Each row in the heat map represents a PSTH of the mean response for a responsive cell-call pair (bin size = 10 ms). Rows are sorted by response magnitude from most inhibitory to most excitatory. *Bottom panels:* Mean traces for excitatory and inhibitory cell-call pairs for each genotype comparing naive females (black) and surrogate females (red). Center and outside lines represent mean and S.E.M., respectively. (B, C) Boxplot of the integrals (area under the curve or AUC) of each inhibitory and excitatory response comparing naive and surrogate WT females. AUC was calculated for the early part of the response (0 – 200 ms relative to stimulus onset) (B) (naive inhibitory: n = 103; surrogate inhibitory: n = 51; naive excitatory: n = 69; surrogate excitatory: n = 67; Mann-Whitney U test with Holm-Bonferroni correction; *\*\*p* < 0.01; n.s. *p* = 0.18) and for the later part of the response (200 – 350 ms relative to stimulus onset) (C) (Mann-Whitney U test with Holm-Bonferroni correction; \**p* < 0.05; *\*\*p* = 0.01). (D, E) Boxplot of the integrals (area under the curve or AUC) of each inhibitory and excitatory response comparing naive and surrogate HET females. AUC was calculated for the early part of the response (0 – 200 ms relative to stimulus onset) (D) (naive inhibitory: n = 80; surrogate inhibitory: n = 51; naive excitatory: n = 116; surrogate excitatory: n = 67; Mann-Whitney U test with Holm-Bonferroni correction; n.s. *p* = 0.41; n.s. *p* = 0.29) and for the later part of the response (200 – 350 ms relative to stimulus onset) (E) (Mann-Whitney U test with Holm-Bonferroni correction; n.s. *p* = 0.75; n.s. *p* = 0.12).

Qualitative comparison of the 2D PSTHs between NaiveWT and SurWT reveals that they are consistent with a shift in abundance and magnitude away from inhibitory responses and towards excitatory responses in SurWT. Comparing traces of the mean excitatory and inhibitory responses between NaiveWT and SurWT, reveals that distinct from the changes in abundance, there are time-dependent differences in the amplitude of responses. To quantify these differences, we integrated the area under the response curve during two time windows: 0 - 200 ms (early) and 200 - 350 ms (late). The longest call stimulus was 125 ms in duration. The integrated area under the mean inhibitory response trace was significantly decreased in SurWT in both early (Figure 6B; NaiveWT: 0.136 [0.13 0.15] Z score*s; SurWT: 0.118 [0.10 0.14] Z score*s; Mann-Whitney U test, *p* < 0.01) and late responses (Figure 6C; NaiveWT 0.055 [0.04 0.08] Z score*s; SurWT: 0.039 [0.01 0.06 Z score*s; Mann-Whitney U test, *p* < 0.05). This pattern is consistent with disinhibition of deep non-PV neurons throughout the response. Interestingly, the integrated area under the mean excitatory response trace was significantly increased in SurWT during the late response (Figure 6C; NaiveWT: 0.010 [0.0 0.03] Z score*s; SurWT: 0.050 [0.03 0.07] Z score*s; Mann-Whitney U test, *p* < 0.01), but not during the early response (Figure 6B; NaiveWT: 0.223 [0.17 0.27] Z score*s; SurWT: 0.181 [0.14 0.24] Z score*s; Mann-Whitney U test, *p* = 0.18). Thus, auditory response plasticity in deep non-PV neurons, potentially reflecting removal of PV neuron-mediated inhibition, resulted in an extension of the excitatory response duration.

In contrast, we found no significant differences in the response integral in deep non-PV neurons between NaiveHET and SurHET during either the early response (Figure 6D; NaiveHET excitatory: 0.203 [0.17 0.23] Z score*s; SurHET excitatory: 0.220 [0.19 0.26] Z score*s; Mann-Whitney U test, *p* = 0.24; NaiveHET inhibitory: 0.145 [0.112 0.16] Z score*s; SurHET inhibitory: 0.143 [0.12 0.15] Z score*s; Mann-Whitney U test, *p* = 0.97) or the late response (Figure 6E; NaiveHET excitatory: 0.013 [0.00 0.02] Z score*s; SurHET excitatory: 0.027 [0.01 0.04] Z score*s; Mann-Whitney U test, *p* = 0.12; NaiveHET inhibitory: 0.055 [0.03 0.06] Z score*s; SurHET inhibitory: 0.052 [0.03 0.06] Z score*s; Mann-Whitney U test, *p* = 0.75).

PV neurons showed a complementary pattern of changes in their firing in the late portion of their response to pup calls. Figure 7A (upper panels) show 2D PSTHs of PV neuron responses in each genotype and maternal experience condition, and the lower panels show mean response traces comparing naive mice (black) to surrogate mice (red). In this case, because nearly all cells showed net excitatory responses, these traces represent the mean of all responses. Qualitatively, these panels reflect changes in PV neuron output in the late response. As above, we quantified these changes by integrating the area under the response curve during the early and late responses. In *Mecp2*^*wt*^, PV neurons showed a significant decrease in their response to calls during the late response (Figure 7C; NaiveWT: 0.089 [0.05 0.14] Z score*s; SurWT: 0.039 [0.01 0.0] Z score*s; Mann-Whitney U test, *p* < 0.01), but not during the early response (Figure 7B; NaiveWT:

**Figure 7:**
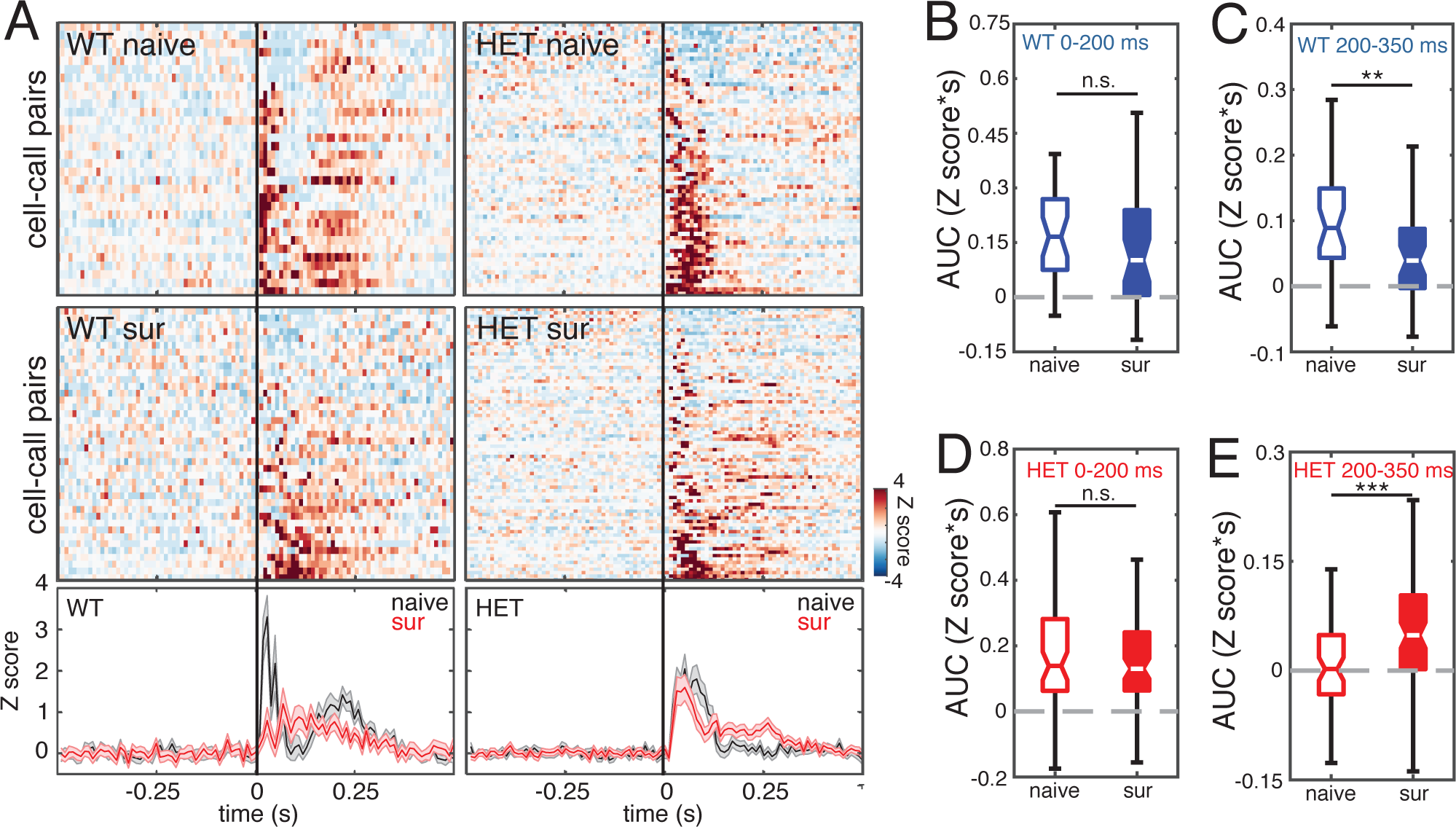
PV neuron output is reduced after maternal experience in WT females and is strengthened after maternal experience in HET females. (A) *Top panels:* Heat maps depicting Z score-normalized responses for all PV cell-call pairs in naive and surrogate WT females and in naive and surrogate HET females. Each row in the heat map represents a PSTH of the mean response for a cell-call pair (bin size = 10 ms). Rows are sorted by response magnitude from least to most excitatory. *Bottom panels:* Mean traces for all cell-call pairs for each genotype comparing naive females (black) and surrogate females (red). Center and outside lines represent mean and S.E.M., respectively. (B, C) Boxplot of the integrals (area under the curve or AUC) of each response comparing naive and surrogate WT females. AUC was calculated for the early part of the response (0 – 200 ms relative to stimulus onset) (B) (naive: n = 32 neurons; surrogate: n = 40 neurons; Mann-Whitney U test; n.s *p* = 0.09) and for the later part of the response (200 – 350 ms relative to stimulus onset) (C) (Mann-Whitney U test; \**\*p* < 0.01). (D, E) Boxplot of the integrals (area under the curve or AUC) of each response comparing naive and surrogate HET females. AUC was calculated for the early part of the response (0 – 200 ms relative to stimulus onset) (D) (naive: n = 64 neurons; surrogate: n = 80 neurons; Mann-Whitney U test; n.s. *p* = 0.56) and for the later part of the response (200 – 350 ms relative to stimulus onset) (E) (Mann-Whitney U test; *\*\*p* < 0.001).

0.166 [0.12 0.16] Z score*s; SurWT: 0.101 [0.06 0.16] Z score*s; Mann-Whitney U test, *p* = 0.09). This is consistent with a model in which the increase in late responses of deep non-PV neurons described above in *Mecp2*^*wt*^ is due to reduced firing in PV neurons.

Suppression of PV-mediated inhibition was not observed in *Mecp2*^*het*^. In fact, SurHET mice showed an increase in the strength of their late response to pup calls relative to NaiveHET (Figure 7E; NaiveHET: 0.002 [-0.02 0.03] Z score*s; SurHET: 0.049 [0.02 0.07] Z score*s; Mann-Whitney U test, *p* < 0.001). Early response strength was not changed (Figure 7D; NaiveHET: 0.139 [0.10 0.22] Z score*s; SurHET: 0.130 [0.10 0.17] Z score*s; Mann-Whitney U test, *p* = 0.56).

### Maternal experience sharpens neuronal tuning for pup calls in Mecp2^wt^

We next assessed whether the auditory response plasticity we observed with maternal experience in deep non-PV neurons of *Mecp2*^*wt*^ coincided with changes in neural selectivity. We found that maternal experience led to stronger, sparser auditory responses and steeper tuning curves for pup vocalizations in *Mecp2*^*wt*^ but not in *Mecp2*^*het*^.

Figure 8A shows a comparison of mean sorted tuning curves for all deep non-PV neurons from NaiveWT (black) and SurWT (red). The use of the term “tuning curve” here is not intended to imply systematic neighbor relationships between stimuli on the x-axis. Auditory responses were significantly stronger in SurWT than in NaiveWT for the most effective call stimuli (Mann-Whitney U test with Holm-Bonferroni correction; \*\**p* < 0.01). Consequently, the tuning curves in SurWT were steeper relative to NaiveWT. The same comparison for NaiveHET and SurHET showed that the mean sorted tuning curves from deep non-PV neurons in these two behavioral conditions (Figure 8B) were indistinguishable (Mann-Whitney U test; *p* > 0.05). We also quantified and compared neural selectivity between naive and surrogate mice by computing lifetime sparseness for each neuron, a quantity that ranges from 0 (equal responses to all stimuli) to 1 (response to only one stimulus). Figure 8C shows that maternal experience leads to a significant increase in mean lifetime sparseness for deep non-PV neurons in *Mecp2*^*wt*^ (NaiveWT: 0.255 [0.20 0.23]; SurWT: 0.381 [0.29 0.48]; Mann-Whitney U test, *p* < 0.001). In contrast, maternal experience did not significantly change lifetime sparseness for deep non-PV neurons in *Mecp2*^*het*^ (NaiveHET: 0.312 [0.26 0.45]; SurHET: 0.374 [0.31 0.40]; Mann-Whitney U test, *p* < 0.05).

**Figure 8:**
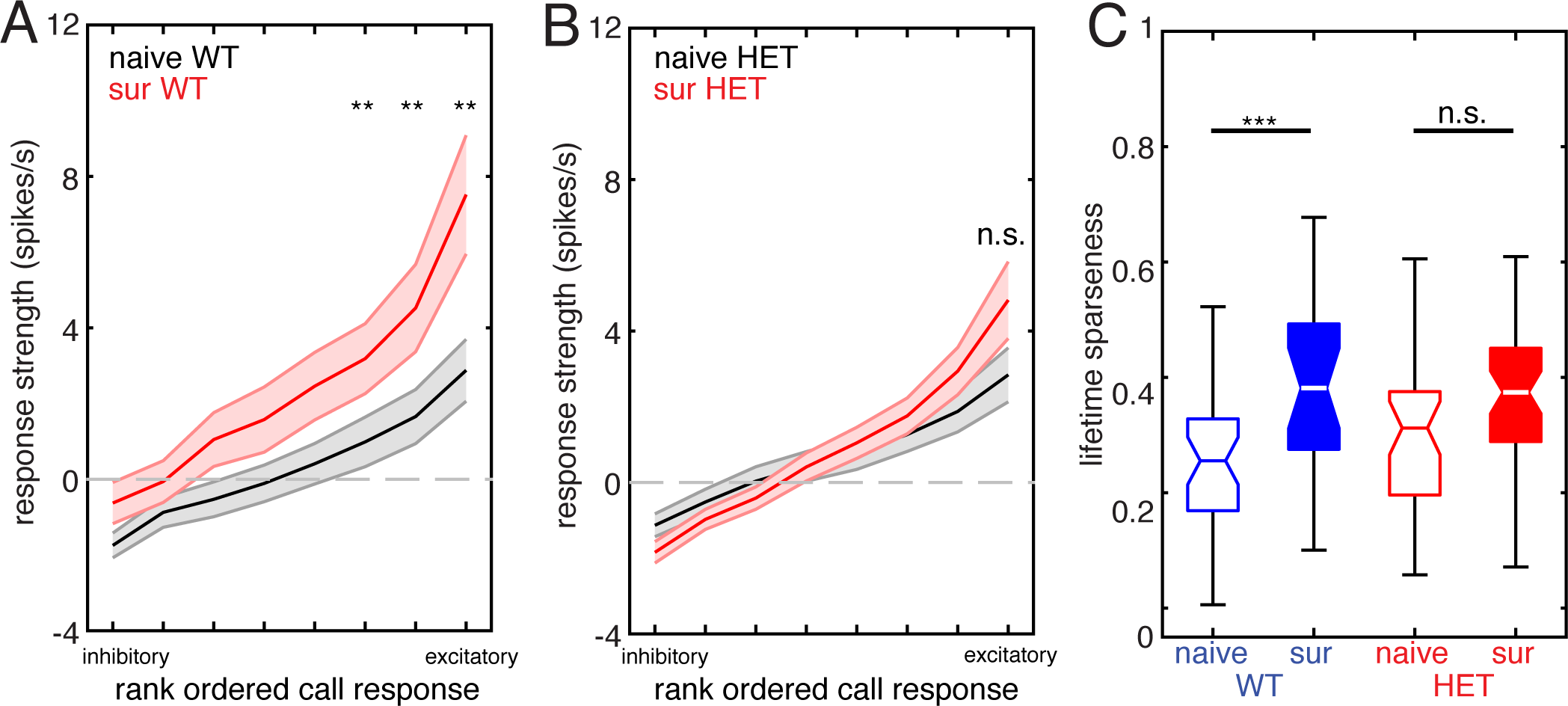
WT females show sharper tuning to calls in deep layer non-PV neurons after maternal experience, but HET females do not. (A) Plot of mean sorted tuning curve for all deep layer non-PV neurons in WT females, comparing naive females (black) and surrogate females (red). Center and outside lines represent mean and S.E.M., respectively. (naive: n = 37 neurons; surrogate: n = 24 neurons; Mann-Whitney U test with Holm-Bonferroni correction: \**\*p* < 0.01). (B) Plot of mean sorted tuning curve for all deep layer non-PV neurons in HET females, comparing naive females (black) and surrogate females (red). Center and outside lines represent mean and S.E.M., respectively. (naive: n = 37 neurons; surrogate: n = 47 neurons; Mann-Whitney U test with Holm-Bonferroni correction: n.s. *p* > 0.05). (C) Boxplot of mean lifetime sparseness computed for each tuning curve from all deep non-PV cells comparing naive females to surrogate females for WT (blue; naive n = 37 neurons and surrogate n = 24 neurons) and HET (red; naive n = 37 neurons and surrogate n = 47 neurons) genotypes (unpaired t-test with Holm-Bonferroni correction; \*\**\*p* < 0.001; n.s. *p* = 0.06).

### Genetic reduction of GAD1 restores maternal behavior and PV neuron plasticity

Elimination of one copy of the gene GAD1 reduces expression of its protein product, the GABA-synthesizing enzyme GAD67, and also ameliorates overexpression of PV and PNNs in the developing visual cortex of male *Mecp2-*null mice (Krishnan et al., 2015). Remarkably, our previous work found that this manipulation of inhibitory networks not only reversed the increase in PV and PNNs triggered by maternal experience in *Mecp2*^*het*^, but it also restored pup retrieval proficiency to wild type levels (Krishnan et al., 2017). Here we show that *Mecp2*^*het*^; *Gad1^het^* exhibit partial reinstatement of maternal experience-induced plasticity of PV neurons and enhanced firing of deep non-PV neurons in response to pup vocalizations.

We made loose patch recordings from the left ‘core’ auditory cortex of 13 female *Mecp2*^*het*^; *Gad1^het^* mice (8 naive and 5 surrogates) for a total of 61 putative non-PV neurons and 23 putative PV neurons. Data collected in response to pup calls were analyzed as in Figures 6 and 7. Figure 9A (upper panels) show 2D PSTHs of non-PV neuron responses recorded in naive and surrogate *Mecp2*^*het*^; *Gad1^het^* mice. Below are mean response traces comparing excitatory responses and inhibitory response for each group. In contrast to the *Mecp2*^*het*^ only mice, the double mutants exhibited increased excitatory responses following maternal experience (Figure 9B; NaiveHET; Gad1^het^: 0.131 [0.11 0.16] Z score*s; SurHET; Gad1^het^: 0.177 [0.13 0.24] Z score*s; Mann-Whitney U test, *p* < 0.05). This change differed from wild type plasticity in that it was only evident in the initial response during the stimulus (the first 120 ms). Also, inhibitory responses in both conditions were small and did not significantly differ across behavioral conditions (Figure 9B; NaiveHET; Gad1^het^: 0.073 [0.06 0.08] Z score*s; SurHET;Gad1^het^: 0.062 [0.04 0.08] Z score*s; Mann-Whitney U test, *p* = 0.24). Nevertheless, recordings from PV neurons in *Mecp2*^*het*^;*Gad1^het^* mice revealed reciprocal maternal experience-dependent changes, with lower spiking output of these inhibitory cells in surrogates in response to calls (Figure 9C,D; NaiveHET;Gad1^het^: 0.079 [0.04 0.11] Z score*s; SurHET;Gad1^het^: 0.036 [0.01 0.07] Z score*s; Mann-Whitney U test, *p* < 0.01). These results are consistent with partial restoration of the pattern of disinhibition observed in wild type mice.

**Figure 9:**
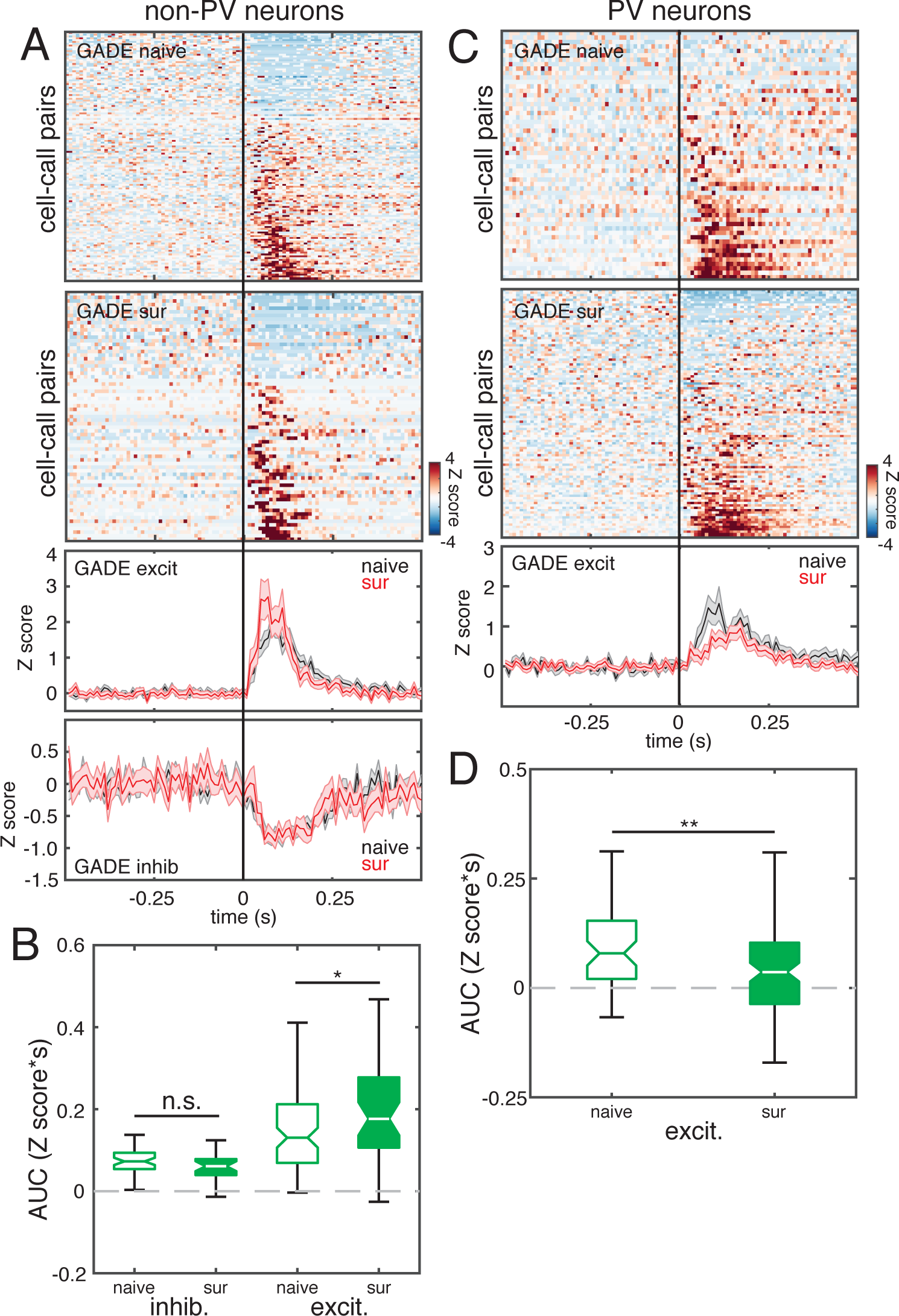
PV neuron plasticity is restored in *Mecp2^het^;Gad1^het^* mice. (A) *Top panels:* Heat maps depicting Z score-normalized responses for all non-PV cell-call pairs in naive and surrogate *Mecp2^het^;Gad1^het^*. Each row in the heat map represents a PSTH of the mean response for a cell-call pair (bin size = 10 ms). Rows are sorted by response magnitude from least to most excitatory. *Bottom panels:* Mean traces for excitatory and inhibitory cell-call pairs for the double mutants comparing naive females (black) and surrogate females (red). Center and outside lines represent mean and S.E.M., respectively. (B) Boxplot of the integrals (area under the curve or AUC) of each response comparing naive and surrogate *Mecp2^het^;Gad1^het^* females. AUC was calculated only for the activity during the stimulus (0 – 120 ms relative to stimulus onset) (naive: n = 39 neurons; surrogate: n = 22 neurons; Mann-Whitney U test; n.s. *p* = 0.24; * *p* < 0.05) (C) Top panels: Organized as in (A) but data are from all PV cell. *Bottom panel:* Mean traces for all cell-call pairs for each genotype comparing naive females (black) and surrogate females (red). Center and outside lines represent mean and S.E.M., respectively. (D) Boxplot of the integrals (area under the curve or AUC) of each response comparing naive and surrogate *Mecp2*^*het*^;*Gad1*^*het*^ females. AUC was calculated only for the activity during the stimulus (0 – 120 ms relative to stimulus onset) (naive: n = 7; surrogate: n = 16; Mann-Whitney U test; ** *p* < 0.01).

## DISCUSSION

Previously, we found that heterozygous female *Mecp2* mutants failed to learn to accurately retrieve pups, and we showed that this was due to a requirement for MECP2 in the auditory cortex at the time of maternal learning (Krishnan et al., 2017). We also found that the absence of *Mecp2* was linked to a rapid and inappropriate maturation of PV networks marked by overexpression of parvalbumin protein and PNNs. Maturation of PV networks is typically associated with the termination of plasticity, such as at the end of developmental critical periods, and the PV neuron population is therefore considered to be a ‘brake’ on plasticity (Levelt and Hubener, 2012; Takesian and Hensch, 2013). Consistent with this framework, in our study the accelerated development of PV and PNNs, and the behavioral deficit, were reversed by genetic deletion of one copy of the gene *Gad1*, which codes for the GABA synthetic enzyme GAD67 (Krishnan et al., 2017).

Here we show that the precipitous maturation of the auditory cortical PV network in *Mecp2*^*het*^ after being exposed to pups for the first time is indeed correlated with impaired plasticity of neuronal responses specifically to vocal signals from pups. The reciprocal modulation of PV and non-PV neurons strongly suggest that PV neurons in wild-type mice exert less inhibition of stimulus-evoked responses of deep non-PV presumptive pyramidal cells. Responses in the deep non-PV neurons are concomitantly sharpened to become more selective for pup calls. In contrast, in *Mecp2*^*het*^, maternal experience-dependent changes in PV neurons are blocked or even reversed. Consequently, no response plasticity and no increase in selectivity was observed in the deep non-PV population of SurHET. Loss of *Mecp2* has been previously shown to abolish or interfere with short- and long-term synaptic plasticity and homeostatic plasticity (e.g. Asaka et al., 2006; Blackman et al., 2012; Na et al., 2013). Notably, deleting one *Gad1* allele, which restores both inhibitory marker levels and behavioral performance, also leads to a partial restoration of PV neuron plasticity and stronger excitatory responses in non-PV neurons. Therefore, taken together our data are consistent with a model in which augmented PV neuron maturation (Patrizi et al., 2019) in *Mecp2*^*het*^ constrains neuronal plasticity in the auditory cortex. We speculate that this plasticity is an important component of early maternal learning. We also propose that altered PV networks in *Mecp2*^*het*^ broadly impede plasticity during episodes of heightened learning in development or adulthood, thereby affecting other behaviors and circuits.

### Effects of maternal experience on auditory cortical circuits in Mecp2^wt^ and Mecp2^het^

Our results show that maternal experience triggers a shift in the balance of excitatory and inhibitory responses to pup vocalizations exhibited by deep non-PV neurons. Specifically, the prevalence and magnitude of call-evoked firing suppression (‘inhibition’) diminished in surrogates, while those of firing increases (‘excitation’) grew. A parsimonious explanation for this shift is that the non-PV neurons become ‘disinhibited’ following maternal experience. Indeed, inhibitory PV neurons appeared to reciprocate with reduced spiking output, inviting us to speculate that they are the source of disinhibition in deep non-PV. We summarize the changes we observed after maternal experience, and we propose an explanatory circuit model, in Figure 10.

**Figure 10:**
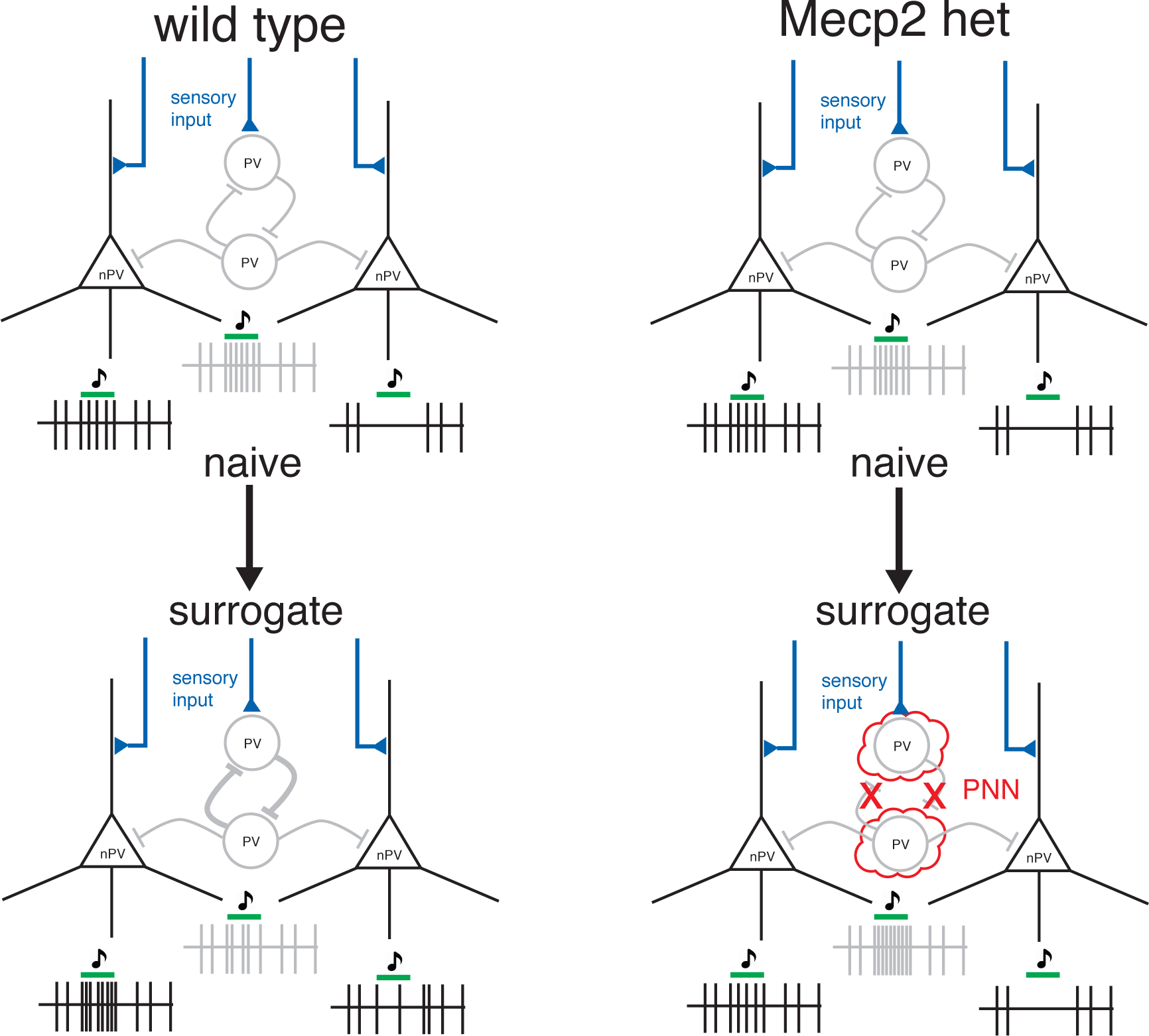
Proposed model for maternal experience-dependent plasticity in wild-type and *Mecp2*^*het*^ mice. In naive wild type mice, we propose that sensory input is received by PV and non-PV (nPV) neurons (in this simplified circuit, non-PV neurons are assumed to be pyramidal neurons), and that spiking output from PV neurons (depicted by the gray spike train) attenuates excitatory responses and enhances inhibitory responses in the non-PV neurons (depicted by the black spike trains). In the transition to surrogacy, PV neuron output is decreased, speculatively depicted here as occurring by stronger self-inhibition of the PV population, disinhibiting the non-PV neurons. Consequently, in non-PV neurons, weakly excitatory responses become stronger, and inhibitory responses become weaker. In *Mecp2*^*het*^, PNNs interfere with synaptic plasticity at connections on PV neurons, thus PV do not disinhibit the non-PV cells.

In naive wild type mice, we propose that sensory input is received by PV and non-PV neurons (Figure 10). In this simplified circuit, non-PV neurons are assumed to be pyramidal neurons. Spiking output from PV neurons (depicted by the gray spike train) attenuates excitatory responses and enhances inhibitory responses in the non-PV neurons (depicted by the black spike trains). In the transition to surrogacy, PV neuron output is decreased, speculatively depicted here as occurring by stronger self-inhibition of the PV population, disinhibiting the non-PV neurons. Consequently, in non-PV neurons, weakly excitatory responses become stronger, and inhibitory responses become weaker. In *Mecp2*^*het*^, PNNs interfere with synaptic plasticity at connections on PV neurons, thus PV do not disinhibit the non-PV cells. We speculate that the lower levels of PNN expression in *Mecp2*^*het*^;*Gad1^het^* mice liberate the PV network from that interference, thereby enabling its plasticity.

Interestingly, the disinhibition we see is stimulus-specific, affecting behaviorally-significant calls but not synthetic sounds. On the other hand, decreased PV spiking was still observed in response to synthetic sounds after maternal experience, suggesting that PV-mediated inhibition sculpts responses to vocalizations, but not simple tones and noise. At this time, it is unclear whether that is related to stimulus meaning or stimulus complexity. Moreover, the disconnect between the changes in PV neurons versus non-PV neurons is consistent with changes in PV neurons leading those in non-PV neurons, as opposed to the other way around.

We also found that some aspects of the surrogacy-induced plasticity that we observed here were most prominent in the late phase of the response to calls. Interestingly, this observation bears similarity to a recent report that secondary auditory cortex (A2) neurons show maternal plasticity in offset responses to frequency-modulated stimuli (Chong et al., 2019). Nevertheless, we are confident that this effect does not bear on our data, because our recordings were made in more dorsal thalamorecipient auditory cortical areas, not in A2.

The changes in inhibition that we found are triggered by maternal experience may partially reflect and synergize with changes in auditory cortical inhibition seen in other studies. For example, Cohen and Mizrahi (2015) observed PV neuron-mediated disinhibition in layer 2/3 of the auditory cortex when anesthetized lactating mothers were exposed to pup odors. Interestingly, this odor-induced form of disinhibition may be distinct from what we found here in deeper layers, which were not examined in that study. Marlin et al (2015) found that the neuropeptide oxytocin, which plays a crucial role in the auditory cortex to establish maternal behavior, leads to acute disinhibition of responses to auditory stimuli, including pup calls. The long-term consequences of early maternal experience beyond weaning however may be increased inhibition, as Galindo-Leon et al (2009) found evidence for in post-weaning mothers.

Analysis of temporal patterns of firing in non-PV and PV neurons revealed that both the disinhibition of non-PV cells and the suppression of firing in PV cells were particularly evident late in the response to vocalizations. These effects synergized to effectively temporally extend responses of non-PV cells to pup calls. Disinhibition of auditory responses may be an important component in shaping neuronal selectivity for vocalizations. Disproportionate enhancement of the average responses to the preferred calls of individual neurons led to steeper tuning curves on average in surrogates. It may also have contributed to a significant increase in lifetime sparseness measured in SurWT compared to NaiveWT. The result of these changes extending response durations and increasing selectivity in deep cells that form the output of the auditory cortex may be a more sensitive representation of calls sent to downstream regions. This may be considered surprising because inhibition is classically believed to play a key role in sharpening selectivity of sensory responses, however recent reexamination of this notion reveals more complexity (Wood et al., 2017). We anticipate this will be a focus of future work.

Nearly all the characteristic changes in PV and non-PV neurons of *Mecp2*^*wt*^ were completely blocked in *Mecp2*^*het*^ (Figure 10). In fact, we found that PV neurons in *Mecp2*^*het*^ showed even greater inhibitory output during the late response to vocalizations. These observations are completely consistent with the accelerated maturation of PV neurons we found in a previous study (Krishnan et al., 2017), which would be expected to increase the strength of inhibitory networks and arrest plasticity. We also used an *Mecp2^flox/flox^* mouse line crossed to *PV-Cre* mice and showed that knockout of *Mecp2* only in PV neurons was sufficient to impair maternal retrieval learning (Krishnan et al., 2017). Collectively, these observations strongly indicate that impaired PV neuron function is a central contributor to maternal behavior deficiencies in *Mecp2*^*het*^.

### Maternal experience reactivates critical period mechanisms

Our data add to a body of evidence suggesting that maternal experience in adult nulliparous or primiparous female mice initiates an episode of heightened learning and plasticity that accesses mechanisms that overlap with those that control developmentally-defined critical periods. It is well-established that developmental critical periods, such as the visual cortical critical period, are regulated by a finely orchestrated sequence of removal, followed by reinstatement, of inhibition (Levelt and Hubener, 2012; Takesian and Hensch, 2013). The termination of these early critical periods is typically accompanied by maturation of intracortical inhibitory networks and intensification of PV expression and PNNs (Levelt and Hubener, 2012; Takesian and Hensch, 2013). We know that these events actively suppress network plasticity because dissolution of PNNs can actually reinstate visual cortical plasticity (Pizzorusso et al., 2002; Pizzorusso et al., 2006). More recent work further establishes that PV expression levels are inversely correlated with both neural plasticity and learning in adult mice (Donato et al., 2013).

Here we showed that in *Mecp2*^*wt*^, maternal experience leads to an enhancement of sensory responses in the auditory cortex that is likely implemented by decreased stimulus-evoked activity in the inhibitory PV neurons. This pattern is shared with critical periods in development, raising the possibility that critical period mechanisms are reactivated in adulthood to facilitate learning. Therefore, early life plasticity mechanisms may be retained and accessed in adulthood, enabling more dynamic control of neural plasticity throughout life.

### Mecp2 mutation interferes with critical periods

Our results are consistent with a broad base of evidence that *Mecp2* specifically plays a role in regulating gene expression programs that govern plasticity during critical periods or other episodes of heightened learning (Picard and Fagiolini, 2019). One early indication of this came from the observation that *Mecp2* regulates activity-dependent expression of BDNF (Chen et al., 2003; Zhou et al., 2006). Moreover, *Mecp2* mutants exhibit accelerated critical periods for visual system development (Krishnan et al., 2015). In our own work, *Mecp2*^*het*^ adult female mice showed similar expression levels of inhibitory markers in the auditory cortex as *Mecp2*^*wt*^ prior to pup exposure (Krishnan et al., 2017). The expression patterns diverged once the mice had acquired maternal experience. Thus, the learning triggered by social experience revealed aspects of *Mecp2*^*het*^ pathology that were previously latent. The idea that *Mecp2* is more essential in developmentally or experientially defined windows is supported by nonlinear developmental trajectory of individuals with Rett syndrome. We propose based on our work that these observations may outline general principles of circuit pathology in *Mecp2* mutation. Specifically, *Mecp2* may act at different times and in different brain regions to regulate learning by modulating the threshold or capacity for circuit plasticity.

## METHODS

### Animals

Adult female mice (7-10 weeks old) were maintained on a 12-h-12-h light-dark cycle (lights on 07:00 h) and received food ad libitum. Genotypes used were CBA/CaJ, *Mecp2*^*het*^ (C57BL/6 background; B6.129P2(C)-*Mecp2^tm1.1Bird^*/J) and *Mecp2*^*wt*^ littermate control. Heterozygous female *Ai32;PV-ires-Cre* mice (*Ai32^+/-^;PV-Cre^+/-^*) were generated by crossing female *Ai32* (B6;129S-*Gt(ROSA)26Sor^tm32(CAG-COP4*H134R/EYFP)Hze^*/J) and male *PV-ires-Cre* (B6;129P2-Pvalbtm1(cre)Arbr/J). The double mutant *Mecp2^het^; Gad1^het^* (*Het;Gad1^het^*) was generated by crossing *Mecp2*^*het*^ females and *Gad1^het^* males. Details on the *Het; Gad1^het^* line can be found in (Krishnan et al., 2017). All procedures were conducted in accordance with the National Institutes of Health’s Guide for the Care and Use of laboratory Animals and approved by the Cold Spring Harbor Laboratory Institutional Animal Care and Use Committee.

### Pup retrieval behavior

Pup retrieval behavior was performed as previously described (Krishnan et al., 2017). Briefly, two virgin female mice (one *Mecp2*^*het*^ and one *Mecp2*^*wt*^) were co-housed with pregnant CBA/CaJ female 3-5 days before birth. Pup retrieval behavior was assessed starting on the day the pups were born (postnatal day 0).

### Surgery for awake recordings

We anesthetized the mice with a mixture of ketamine (100 mg/mL) and xylazine (20 mg/mL) and maintained with isoflurane. We affixed a head bar to the skull above the cerebellum using Rely X cement (3M) and methyl methacrylate-based dental cement (TEETS). For additional stability, we secured 5 machine screws (Amazon Supply) to the skull prior to the application of cement. Animals were allowed to recover >24 hours before used for recordings. For surrogate mice, surgery was performed immediately after behavioral assessment on day 5, and recordings were obtained between days 7 and 11.

### Electrophysiology

On the day of recording, mice were re-anesthetized with isoflurane and a craniotomy was made to expose the left hemisphere of the auditory cortex. Mice were then head-fixed via the attached head bar over a Styrofoam ball that was suspended above the air table. The Styrofoam ball allowed mice to walk and run in one dimension (forward-reverse).

Stimuli were presented via ED1 Electrostatic Speaker Driver and ES1 Electrostatic Speaker (TDT) positioned 4 inches directly in front of the animal. Experiments were performed in a sound attenuation chamber (Industrial Acoustics) at 65 dB SPL RMS measured at the animal’s head using a sound level meter (Extech) with A-weighting by normalizing to the SPL of an 8 kHz reference tone. The speaker used has +/− 11 dB output between 4 – 110 kHz. Stimuli consisted of broadband noise, four logarithmically-spaced tones ranging between 4 and 32kHz, ultrasound noise bandpassed between 40 and 60 kHz, and 8 natural pup calls recorded from 2-4 postnatal day wild type CBA/CaJ mouse pups, digitized at a sampling rate of 195.3 kHz. All stimuli were low pass filtered and amplified between the ADC and the speaker driver at 100 kHz with a custom analog filter and preamp (Kiwa Electronics). Pup calls were recorded from pups isolated in a cage inside a sound attenuation chamber (Industrial Acoustics) with an ultrasound microphone (Avisoft) positioned approximately 30 cm above the pup.

Single units were blindly recorded *in vivo* by the ‘loose patch’ technique using borosilicate glass micropipettes (7-30 MΩ) tip-filled with intracellular solution (mM: 125 potassium gluconate, 10 potassium chloride, 2 magnesium chloride and 10 HEPES, pH 7.2). Spike activity was recorded using BA-03X bridge amplifier (npi Electronic Instruments), low-pass filtered at 3 kHz, digitized at 10 kHz, and acquired using Spike2 (Cambridge Electronic Design). The depth of the recorded cell was measured by a hydraulic manipulator (Siskiyou MX610; Siskiyou Corporation).

### Photostimulation-assisted identification of PV cells

In addition to the head-bar implant, naive *Ai32^+/-^;PV-Cre^+/-^* mice were implanted with an optic fiber (Ø105 um, NA 0.22; ThorLabs) pigtailed to a ceramic ferrule (Ø128 um hole size; Thorlabs), stabilized with TEETS dental cement, and with the tip inserted into the left hemisphere of the auditory cortex. Stimuli consisted of trains of ten 10-ms pulses of blue light (473 nm; OEM Laser) delivered at 2 Hz, with power of 2.5-4.5 mW measured at the tip of optic fiber.

### Data analysis

All data were manually spiked sorted into single-unit spike trains with Spike2 (CED) and subsequently analyzed using custom software written in Matlab (MathWorks).

For photo identification of PV neurons (Lima et al, 2009), we computed the median latency over all trials to the first spike following light onset (trials with latency >50ms were set at 50 ms), and the reliability as the proportion of trials where the cell produced at least one spike within 50 ms of light onset. Identified PV neurons exhibited latencies of a few ms and reliability close to 1. Peristimulus time histograms (PSTHs) in Figure 1A depict spike probability in each bin as the number of spikes in all trials divided by the total number of trials. We identified the peak and trough of the mean spike waveform (from the spikes occupying the quartile from 70 – 90 percentile amplitude) for every neuron reported in this study. We computed the interval between them and the ratio of the amplitudes of the trough and the peak. Based on our photo identification results, neurons with waveforms that had an interval of 0.5 ms or less and a trough/peak amplitude ratio of >0.5 were designated as putative PV neurons. All other neurons were designated as putative non-PV neurons. Depth was recorded for all neurons, and neurons at a depth of <500 µm were designated “superficial layer”, and neurons at a depth of >500 µm were designated “deep layer”. Data from neurons deeper than 1100 µm were discarded.

For the analysis in Figures 6, 7 and 9, we constructed a PSTH (10 ms bin size) of the mean spiking response across trials to each stimulus for each neuron. The bin values for the mean response to all stimuli were transformed to Z scores. A bootstrap procedure was used to identify all individual cell-call pairs with a significant response to the call. Briefly, we randomly sampled 15 bins from all histograms for a given cell 10,000 times to create a null distribution, and we compared the mean bin value during the 150 ms after the stimulus. Responses that fell above the top 2.5%, or below the bottom 2.5%, of this distribution were considered significantly excitatory or inhibitory respectively. Only significant responses were included in this analysis.

Response strength was computed as the mean firing rate during the first 350 ms after stimulus onset, minus the mean firing rate during the preceding 3 s (which was also used to compute the reported spontaneous firing rate for each cell). Lifetime sparseness was calculated as follows (Meeks et al., 2010):

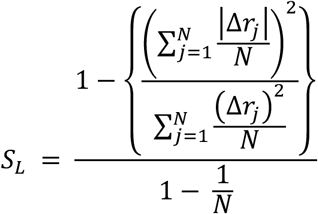

*N* is the number of different stimuli presented to the neuron, and Δ*r*_*j*_ is the change in firing rate of the neuron in response to the *j*th stimulus.

## Acknowledgements

We wish to thank J. Hall, R. Prosser, and R. Liu for helpful comments and discussion. This work was supported by grants to SDS from the Simons Foundation Autism Research Initiative (SFARI), the Feil Family Foundation, and the National Institute of Mental Health (R01MH106656), to ZJH from the National Institute of Mental Health (R01MH102616), and to KK from National Alliance for Research on Schizophrenia and Depression Young Investigator Grant, and the Brain and Behavior Research Foundation, and by International Rett Syndrome Foundation Postdoctoral Fellowships to BYL and KK. Additional support came from laboratory startup funds to KK from University of Tennessee-Knoxville.

